# The genomic signatures of natural selection in admixed human populations

**DOI:** 10.1101/2021.09.01.458503

**Authors:** Sebastian Cuadros Espinoza, Guillaume Laval, Lluis Quintana-Murci, Etienne Patin

**Author notes:** Corresponding authors (S.C.-E.), (E.P.).

## Abstract

Admixture has been a pervasive phenomenon in human history, shaping extensively the patterns of population genetic diversity. There is increasing evidence to suggest that admixture can also facilitate genetic adaptation to local environments, i.e., admixed populations acquire beneficial mutations from source populations, a process that we refer to as *adaptive admixture*. However, the role of adaptive admixture in human evolution and the power to detect it are poorly characterized. Here, we use extensive computer simulations to evaluate the power of several neutrality statistics to detect natural selection in the admixed population, accounting for background selection and assuming different admixture scenarios. We show that two statistics based on admixture proportions, *F*_adm_ and LAD, show high power to detect mutations that are beneficial in the admixed population, whereas iHS and *F*_ST_ falsely detect neutral mutations that have been selected in the source populations only. By combining *F*_adm_ and LAD into a single statistic, we scanned the genomes of 15 worldwide, admixed populations for signatures of adaptive admixture. We confirm that lactase persistence and resistance to malaria have been under adaptive admixture in West Africa and in Madagascar, North Africa and South Asia, respectively. Our approach also uncovers new cases of adaptive admixture, including the *APOL1*/*MYH9* locus in the Fulani nomads and *PKN2* in East Indonesians, involved in resistance to infection and metabolism, respectively. Collectively, our study provides new evidence that adaptive admixture has occurred in multiple human populations, whose genetic history is characterized by periods of isolation and spatial expansions resulting in increased gene flow.

**Author summary:** Adaptive introgression, i.e., the acquisition of adaptive traits through hybridization with another species, is a well-documented phenomenon in evolution. Conversely, adaptive admixture, i.e., the acquisition of adaptive traits through admixture between populations of the same species, is poorly described. In this study, we evaluate the importance of adaptive admixture in human recent evolutionary history. We first determine the expected signatures of adaptive admixture on patterns of genomic diversity, using realistic simulations. We then identify the methods that are the most powerful to detect such molecular signatures. Finally, by using the methods identified as the most powerful, we search for cases of adaptive admixture in the genomes of 15 admixed populations from around the globe. We find evidence that adaptive admixture has occurred in several populations from Northeast Africa, Southeast Asia and Oceania. This study suggests that admixture has indeed facilitated human genetic adaptation, particularly at genes involved in metabolism and resistance against pathogens.

## Introduction

Over the last two decades, the search for molecular signatures of natural selection in the human genome has played an integral part in understanding human evolution [1–6]. Genome scans for local adaptation have shed light on the environmental pressures that populations have faced for the last 100,000 years, including reduced exposure to sunlight, altitude-related hypoxia, new nutritional resources or exposure to local pathogens. Candidate genes for local genetic adaptation have been identified based on expected signatures of positive selection, such as extended haplotype homozygosity or strong differences in allele frequencies between geographically diverse populations. In doing so, selection studies have implicitly assumed that advantageous variation occurred in a single population that has remained isolated from other populations since their separation. Yet, recent ancient and modern genomics studies have demonstrated that the last millennia of human history have been characterized by large-scale spatial expansions, followed by extensive admixture [1,7,8]. This suggests that expanding groups, as they migrated, met and admixed with the local populations they encountered, may have inherited alleles that increased their fitness in the newly settled environments. This phenomenon, which we refer here as *adaptive admixture*, may have played an important role in human evolution.

While an increasing number of studies have revealed how admixture with ancient hominins, such as Neanderthals or Denisovans, facilitated modern human adaptation [9], the adaptive nature of admixture between modern humans remains largely unexplored. Nonetheless, there is suggestive evidence that adaptive admixture has indeed occurred in humans, as several studies have reported candidate loci for positive selection in admixed populations [10–30]. A striking example is the Duffy-null *FY*B^ES^* allele, which confers protection against *Plasmodium vivax* malaria [31,32]. Signals of selection since admixture have been detected at the locus in diverse, African-descent, admixed populations from Madagascar, Cabo Verde, Sudan and Pakistan [12,19,20,23,25], suggesting strong, ongoing selection owing to *vivax* malaria in these regions. A variety of methods has been used to detect the signatures of adaptive admixture, relying on classic neutrality statistics, such as iHS or *F*_ST_, and deviations from allele frequencies [12,33] or ancestry proportions [11,14,17,20,21,25–29] expected under admixture and neutrality. However, the power of these neutrality statistics to detect adaptive admixture is currently unknown. More worrying, it has been suggested that selection in source populations can confound signals of adaptive admixture [21,22] and, conversely, admixture may obscure signals of positive selection occurring in source populations [34]. Finally, the extent to which adaptive admixture has contributed to human genetic adaptation remains poorly known, as reported signals of adaptive admixture are limited to few populations, relative to the large number of admixture events reported in humans [1,7,8].

In this study, we estimated and compared the power of various neutrality statistics to detect adaptive admixture, through computer simulations under different admixture and selection scenarios. We then used the most powerful statistics to scan the genomes of 15 different admixed human populations from around the world and detect candidate loci for adaptive admixture. In doing so, we confirm several, iconic signals and identify new cases of ongoing positive selection since admixture, which highlight pathogens as key drivers of recent genetic adaptation in humans.

## Results

### Power estimation under different models of admixture with selection

To estimate the power to detect adaptive admixture, we performed extensive forward-in-time simulations of a population that originates from admixture between two source populations. We introduced a beneficial mutation in one of the source populations, with a varying selection coefficient (Methods). We considered three different scenarios of admixture with selection (Fig 1A). *Scenario 1* corresponds to adaptive admixture, where the admixed population inherits adaptive variation from one of its source populations: the beneficial mutation is under positive selection in the source population, is transmitted to the admixed population and remains beneficial – with the same selection coefficient – in the admixed population. In *scenario 2*, the new beneficial allele is under positive selection in the source population, is transmitted to the admixed population and becomes neutral in the admixed population only. We simulated this scenario to verify if some neutrality statistics wrongly support positive selection in the admixed population because of a residual signal inherited from the source population. At the same time, this scenario is also useful to evaluate the power to detect residual signals of positive selection in the admixed population, as a means to study the past evolutionary history of source populations that no longer exist in an unadmixed form [34–36]. Finally, in *scenario 3*, a neutral mutation in the source population becomes beneficial in the admixed population only, at the time of admixture. This case is used to determine how neutrality statistics behave when natural selection operates since admixture on standing neutral variation.

**Fig 1.**
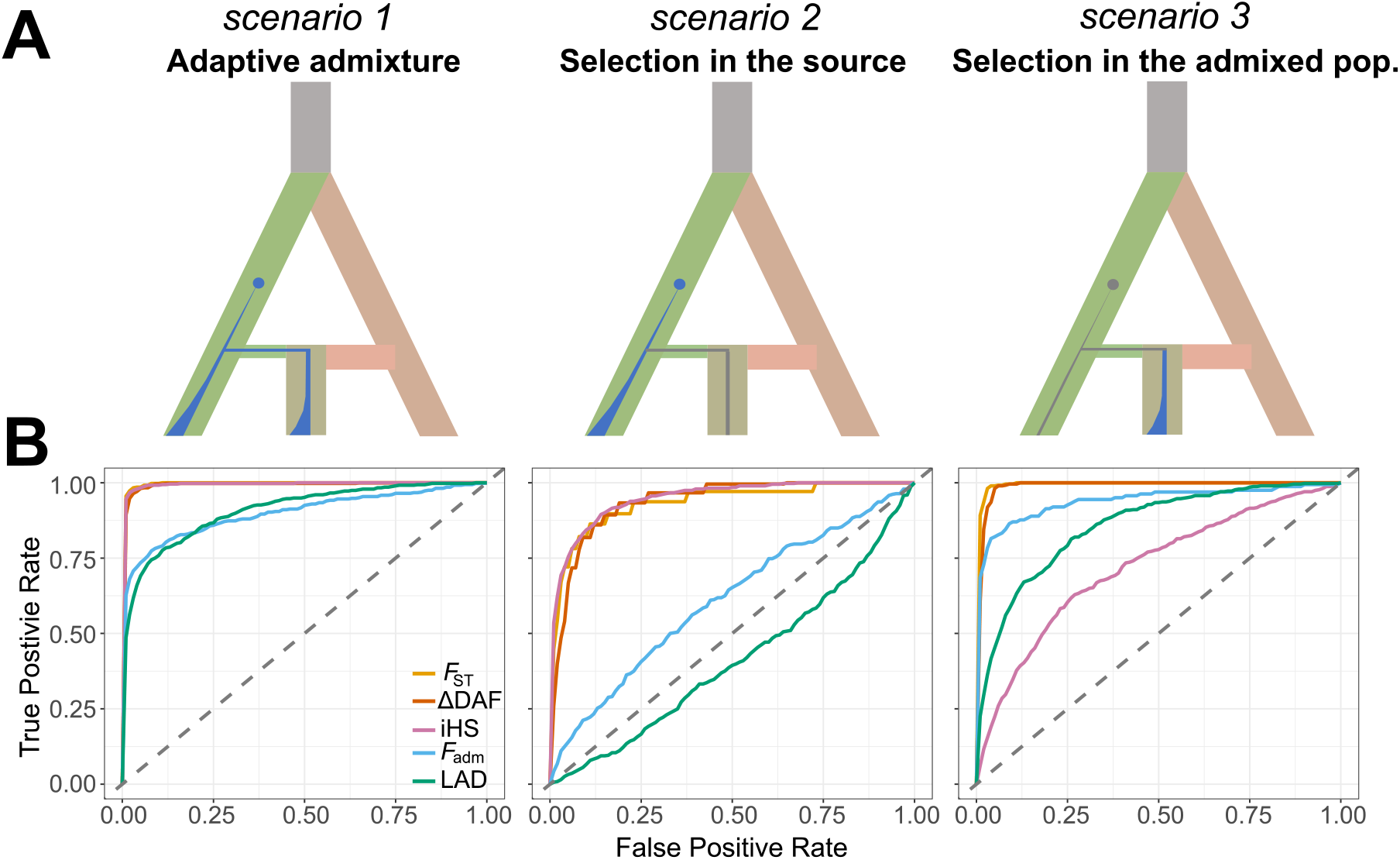
Performance of neutrality statistics under different scenarios of admixture with selection. A. Explored admixture with selection scenarios, from left to right: adaptive admixture, ancestral selection and post-admixture selection on standing variation. The blue and gray points indicate the appearance of a new beneficial and neutral mutation, respectively. The blue and gray areas indicate changes in frequency of the beneficial and neutral mutation, respectively. B. Receiver operating characteristic (ROC) curves comparing the power of classic neutrality statistics *F_ST_*, iHS, ΔDAF and the admixture-specific statistics *F*_adm_ and LAD, across the 3 explored scenarios. Selection coefficient was fixed to *s* = 0.05, to highlight the differences between statistics and between models (see S1 Fig for different selection coefficient values). False positive rate (FPR) is the fraction of simulated neutral sites that are incorrectly detected as adaptive, and true positive rate (TPR) is the fraction of simulated adaptive mutations that are correctly detected as under selection.

We evaluated the performance, under each scenario, of three classic neutrality statistics, *F*_ST_, ΔDAF and iHS, as well as two statistics that are specifically designed to detect selection in an admixed population: *F*_adm_, which is proportional to the squared difference between the observed and the expected allele frequency in the admixed population [12,37,38] and LAD, the difference between the admixture proportion at the locus and its genome-wide average [29], estimated based on local ancestry inference (LAI) by RFMix (Methods) [39]. Receiver operating characteristic (ROC) curves indicate that both the classic neutrality statistics and *F*_adm_ and LAD are powerful to detect adaptive admixture (*scenario 1*) when the selection coefficient *s* = 0.05 (>70% detection power for a false positive rate (FPR) of 5%; Fig 1B, S1 Fig and S2 Fig), in agreement with a previous study [22]. Nevertheless, the power of *F*_ST_, ΔDAF and iHS is also high when the mutation is beneficial in the source population and is no longer selected in the admixed population (*scenario 2*), indicating that these statistics wrongly detect selection in the source population as selection in the admixed population. In contrast, *F*_adm_ and LAD detect adaptive admixture specifically, as their power under *scenario 2* is low or nil (Fig 1B). Of note, our simulations also imply that the power of classic statistics is substantial when using the admixed population as a means to detect selection in the source populations (>65% detection power when *s* = 0.05 and FPR = 5%) [34–36]. Finally, LAD and iHS showed a reduced power to detect selection in the admixed population when the mutation is neutral in the source populations (*scenario 3*, Fig 1A), relative to the adaptive admixture case (*scenario 1*). This may stem from the fact that, under *scenario 3*, the beneficial mutation has been selected for less generations than in *scenario 1*, resulting in a weaker signal for classic statistics. Furthermore, this scenario resembles a scenario of selection on standing variation, where the adaptive mutation may be present on several haplotypes, making it harder to detect [40].

Collectively, our simulations indicate that *F*_adm_ and LAD are the only studied statistics that have substantial power to detect specifically strong, ongoing selection in the admixed population and have more power to detect adaptive admixture than post-admixture selection on standing variation. Because our objective is to detect the signatures of positive selection in the admixed population, and not in the source populations, we based all subsequent analyses on the *F*_adm_ and LAD statistics.

### Effects of the study design

We investigated how sample size and the choice of source populations affect the power of *F*_adm_ and LAD to detect adaptive admixture signals (Methods). We explored sample sizes ranging from *n* = 20 to *n* = 500, for both the admixed and the source populations. We found that *n* = 100 already provides optimal power, because the variance of neutrality statistics is virtually unchanged when *n* ≥ 100 (Fig 2A and S3A Fig). Conversely, we found that when *n <* 50, sampling error increases the variance of *F*_adm_ and LAD null distributions, by as much as 5 times, and ultimately decreases detection power by up to 40% (FPR = 5%). Interestingly, LAD detection power is not affected by the sample size of the source populations, even when *n* = 20 (S3B Fig). Consistently, RFMix accuracy was shown to be only minimally reduced when the sample size of reference panels is as small as *n* = 3 [39], as it uses both source and admixed individuals for LAI.

**Fig 2.**
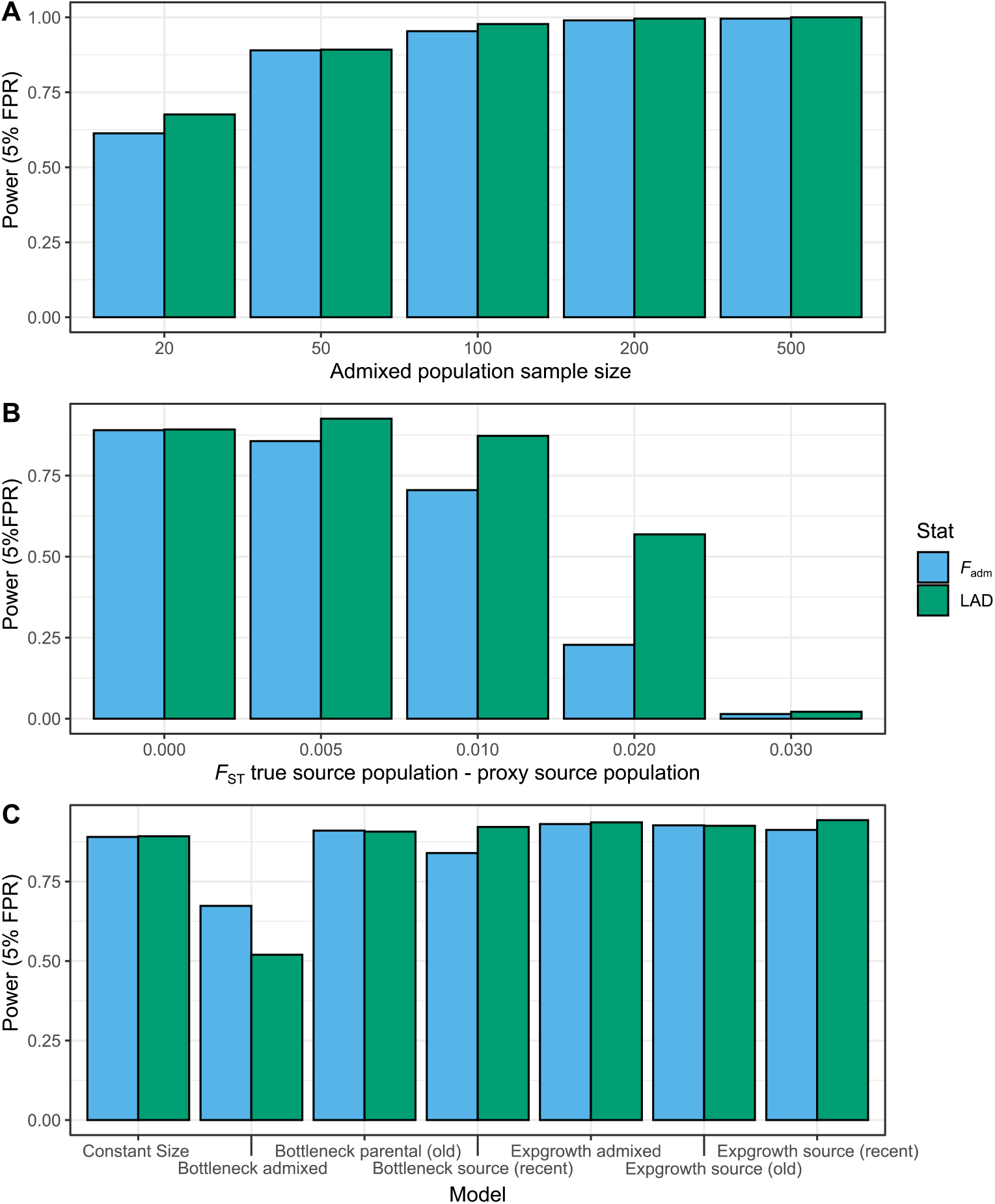
Effects of study design on the power to detect adaptive admixture. Effects of (A) the sample size of the admixed population, (B) the use of source population proxies and (C) non-stationary demography on the detection power of *F*_adm_ and LAD, at a fixed FPR = 5%.

Because obtaining genotype data for the true source populations of an admixed population is difficult, if not impossible, population geneticists often use related, present-day populations as proxies, which may lead to false adaptive admixture signals [21,22]. We explored how detection power is affected by the genetic distance between the true source population and a related population, used as a proxy for *F*_adm_ and LAD computations (Methods). We observed a difference in performance between *F*_adm_ and LAD, the latter being more robust to the use of a proxy (Fig 2B). LAD maintains similar detection power even if the divergence between the true and proxy populations (measured by *F*_ST_) is 0.01, whereas power decreases by 25% for *F*_adm_. This difference in power may stem from the nature of the two statistics. In the case of *F*_adm_, the ancestral allele frequency is directly estimated from the allele frequencies observed in the proxy, and these frequencies are decreasingly correlated with those in the true source population, as their divergence increases. On the other hand, LAD is derived from LAI by RFMix, which was shown to be robust to the use of proxy reference populations [39].

We identified a potentially problematic scenario for both *F*_adm_ and LAD involving population proxies: when the selection event occurs specifically in the proxy source population (i.e., the mutation is not selected in both the true source and the admixed populations), spurious deviations in local ancestry and in allele frequencies were observed in the admixed population. More specifically, this generates an excess of local ancestry from the other source population and expected allele frequencies higher than those observed in the admixed population (S4A and S4B Fig). We found that this scenario produces weaker LAD values (i.e., lower detection power) but stronger *F*_adm_ values (i.e., higher detection power), relative to an adaptive admixture event (S4C Fig). To remediate this, we performed a selection scan in the proxy population using a single-population statistic, iHS, and excluded the top 1% values. In doing so, we managed to exclude approximately 90% of the outlier values of *F*_adm_ and LAD generated by this scenario. More importantly, because there is no correlation, in the case of adaptive admixture, between iHS in the source population and *F*_adm_ or LAD, none of the outlier values generated by a true adaptive admixture event were excluded by this analysis step (S4D and S4E Fig).

### Effects of the admixture model and non-stationary demography

Several studies have shown that admixture in humans has often involved multiple admixture pulses from two or more source populations [8,41–46]. We thus estimated the detection performance of *F*_adm_ and LAD under admixture models that are more complex than the single admixture pulse. We found that the power to detect adaptive admixture is only moderately reduced under a two-pulse admixture model or a constant, continuous admixture model: the true positive rate (TPR) decreases by <11% at a FPR = 5%, relative to the single pulse model (S5A Fig). This suggests that our power estimations are valid for a variety of admixture models.

Assuming a single-pulse admixture model, we then explored how detection performance is impacted by key parameters of the adaptive admixture model, including the admixture time, the admixture proportions and the strength of selection (S1 Table). As expected, we found that detection power is high only when positive selection is very strong; the TPR is up to 94% and 27% when *s* = 0.05 and 0.01, respectively (FPR = 5%; Fig 3 and S6-S10 Figs). Power is also determined by the admixture time *T*_adm_, as it affects the duration of selection; the TPR is up to 94% and 21% when *T*_adm_ ≥ 70 and ≤ 20 generations, respectively. Interestingly, we observed the opposite trend for admixture proportions: the higher the admixture proportion *α* (from the source population where the selected mutation appeared), the lower the detection power. Power decreases particularly when *α >* 0.65, probably because of a threshold effect: if the beneficial allele is at high frequency and e.g., *α =* 0.9, there is little room for the observed allele frequency or local ancestry to deviate from its expectation, making it hard to detect.

**Fig 3.**
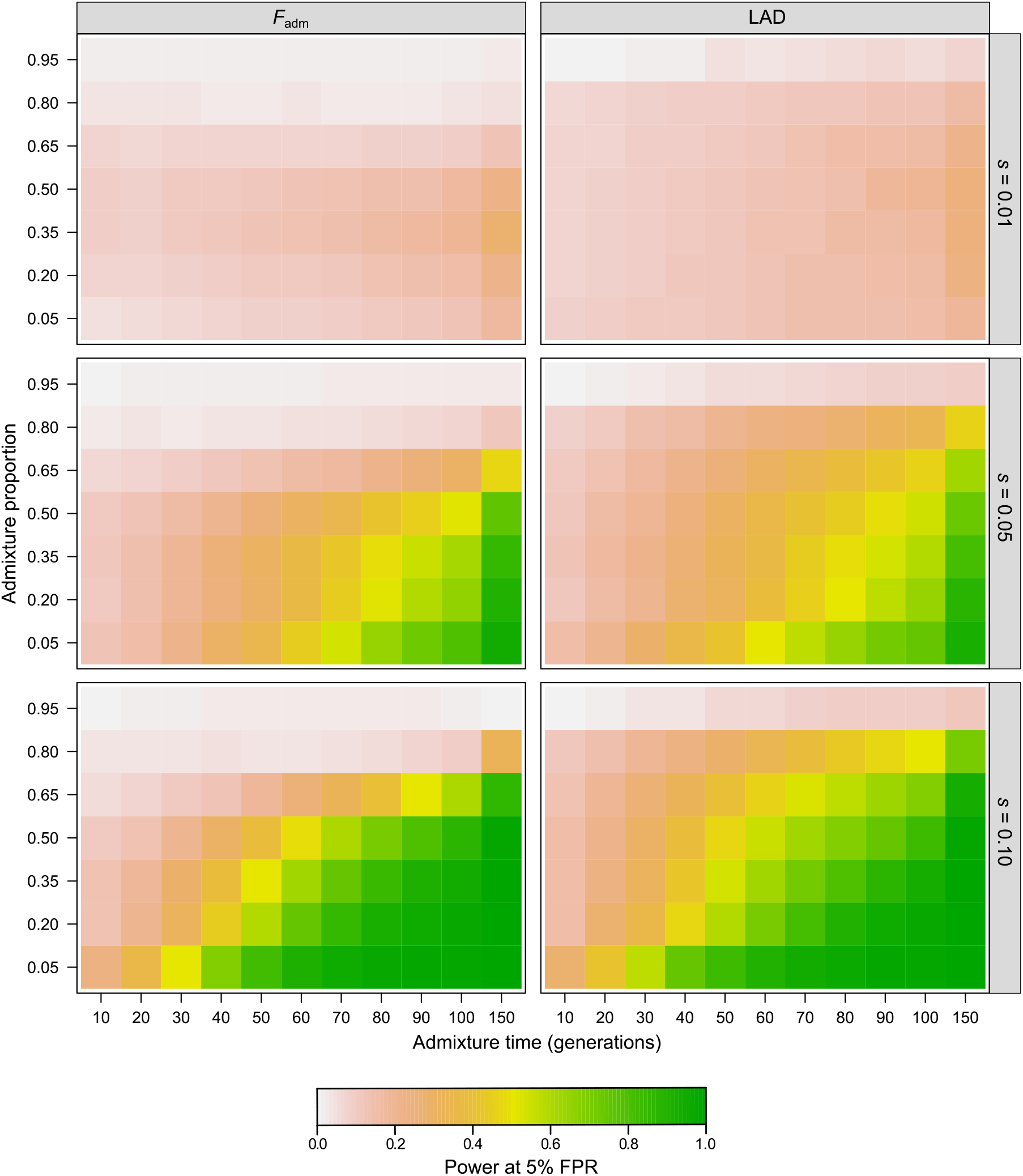
Effects of model parameters on the power to detect adaptive admixture. Color represents average detection power for a fixed FPR = 5% across parameter combinations. The effects of other parameters, such as effective population size and divergence time (S1 Table), are shown in S6-S10 Figs.

We also estimated power under scenarios where demography deviates from a constant population size model. Indeed, demographic events, such as bottlenecks, have been shown to alter the performance of several neutrality statistics [47–53]. We simulated 5 demographic scenarios, including 10-fold bottlenecks and 5% growth rate expansions in either the admixed or the source populations (Methods). We found that detection power is minimally affected under all expansion models (TPR decrease of 5% at a FPR = 5%; Fig 2C). In contrast, detection power is reduced by as much as 50% under the scenario where a 10-fold bottleneck is introduced in the admixed population, relative to the stationary model. This is probably explained by the increased variance of *F*_adm_ and LAD null distributions under this scenario (S5B Fig). Finally, we found that detection power of both *F*_adm_ and LAD is minimally affected when the 10-fold bottleneck is introduced in the source populations, even when it is introduced few generations before the admixture pulse (TPR decrease of 5% at a FPR = 5%; Fig 2C), suggesting that both statistics are robust to increased genetic drift occurring in the source populations.

### Empirical detection of adaptive admixture in humans

We next sought to detect candidate genes for adaptive admixture in humans, by scanning with *F*_adm_ and LAD statistics the genome of 15 worldwide populations (S2 Table) that have experienced at least one admixture event in the last 5,000 years (i.e., the upper detection limit set for accurate local ancestry inference; see [54]). To improve detection power and facilitate candidate prioritization, we combined the empirical *P*-values of both statistics with Fisher’s method [55], used here as a combined test for positive selection since admixture. We confirmed with simulations that the Fisher’s score follows a *χ*^2^ distribution with 4 degrees of freedom under the null hypothesis of absence of positive selection (Fig 4A). Consistently, we found that *F*_adm_ and LAD statistics are not correlated under the null hypothesis (Spearman’s coefficient = 0.03), whereas they are correlated under adaptive admixture (Spearman’s coefficient = 0.96). Furthermore, we found that Fisher’s method increases detection power under unfavourable scenarios, relative to each individual statistic (Fig 4B). In particular, the Fisher’s method improves power when the admixed population experienced a 10-fold bottleneck, when admixture is recent (*T*_adm_ =10 generations) and when using a proxy population that experienced strong drift (*F*_ST_ with the true source population = 0.02). Due to the current lack of knowledge about the recent demographic history of the studied populations, which could increase FPR (S5B Fig), we applied a conservative Bonferroni correction on Fisher’s *P*-values, considering the number of RFMix genomic windows as the effective number of tests (all SNPs within a given window have the same value for LAD). This yielded a *P*-value threshold of approximately *P =* 3.5×10^−6^ (S3 Table). Finally, we verified that the empirical distribution of Fisher’s *P*-values is uniform in all studied populations and found an excess of low *P*-values for several populations (S11 Fig), suggesting that adaptive admixture has occurred in these groups.

**Fig 4.**
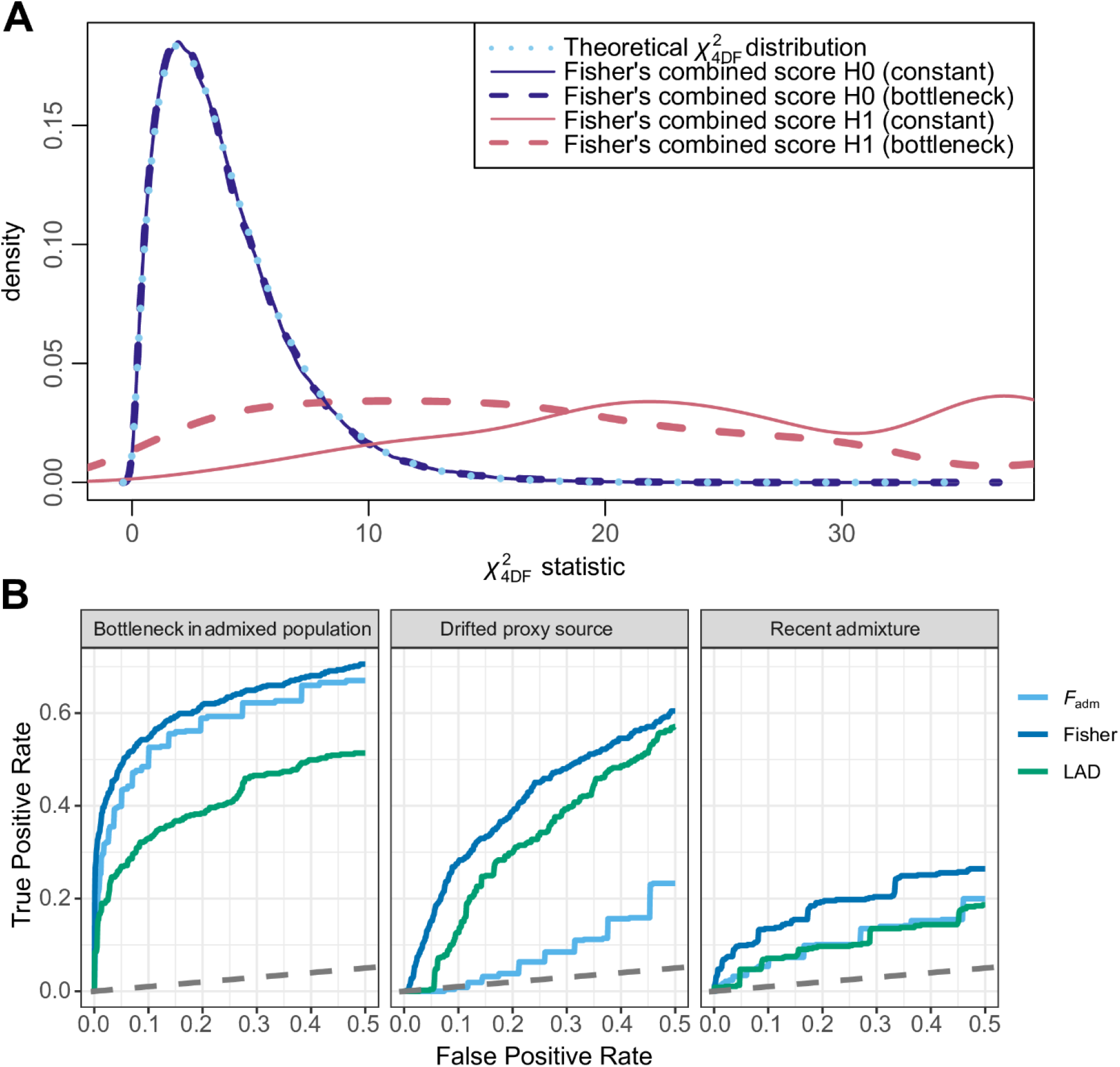
Performance of the Fisher’s method to detect adaptive admixture. (A) Distributions of the combined Fisher’s score under the null hypothesis of no positive selection (H0, blue lines) and under adaptive admixture (H1, pink lines), compared to the theoretical χ^2^ distribution with 4 degrees of freedom (dotted light blue line). Solid and dashed lines indicate distributions under a constant population size and a 10-fold bottleneck in the admixed population. (B) ROC for *F*_adm_, LAD and the combined Fisher’s score under unfavourable scenarios for detecting adaptive admixture: a 10-fold bottleneck introduced in the admixed population, the use of a source population proxy having experienced strong drift (*F*_ST_ between the true source and proxy populations of 0.02) and recent admixture (*T*_adm_ < 30 generations).

Our genome scans indeed identified a number of previously reported signals of adaptive admixture. Among these, we found the *HLA* class II locus in Bantu-speaking populations from Gabon [21] (Fig 5A and 5C; top ranking SNP identified in *HLA-DPA1*; *P =* 7.9×10^−8^; expected frequency of 0.33 *vs.* observed frequency of 0.70), the lactase persistence-associated *LCT/MCM6* locus in the Fulani nomads of Burkina Faso [52] (Fig 6A; top ranked SNP identified in *CCNT2*; *P =* 1.1×10^−6^; expected frequency of 0.12 *vs.* observed frequency of 0.47), and the *ACKR1* gene (previously referred as *DARC*) in African-descent populations from Madagascar, the Sahel and Pakistan [19,20,25] (Fig 5B, 5D and S12 Fig). Furthermore, for the latter locus, the top-ranking variant is *rs12075* in the Malagasy (*P =* 3.4×10^−9^; expected frequency 0.45 *vs.* observed frequency of 0.93), as previously found [20]. This variant, also known as the Duffy-null *FY*B^ES^* allele, confers resistance against *Plasmodium vivax* infection in sub-Saharan Africans [31,32]. Together, these results confirm that our conservative genome scans can recover strong, well-documented signals of adaptive admixture.

**Fig 5.**
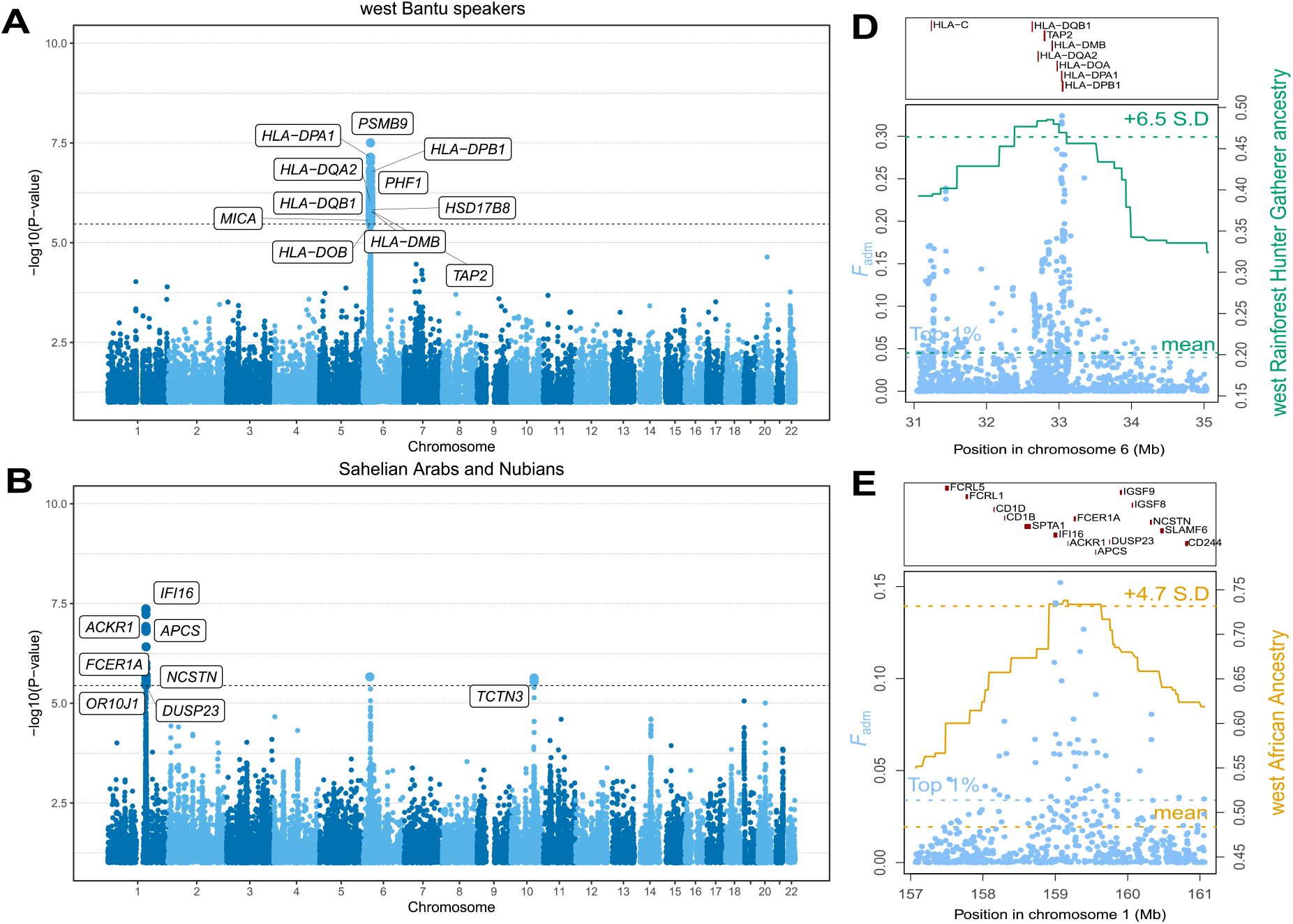
Iconic genomic signals of adaptive admixture. Genome-wide signals of adaptive admixture in (A) Bantu-speaking populations from Gabon and (B) Sahelian Arabs and Nubians. Highlighted blue points indicate variants that passed the Bonferroni significance threshold (shown by a horizontal dotted line). Gene labels were attributed based on the gene with the highest V2G score within 250-kb of the candidate variant. (C-D) Local adaptive admixture signatures for (C) the *HLA* region in Bantu-speaking populations from Gabon and the *ACKR1* region in Sahelian Arabs and Nubians. Light blue points indicate *F*_adm_ values for individual variants. The green and gold solid lines indicate average local ancestry from African rainforest hunter-gatherers and West Africans respectively.

**Fig 6.**
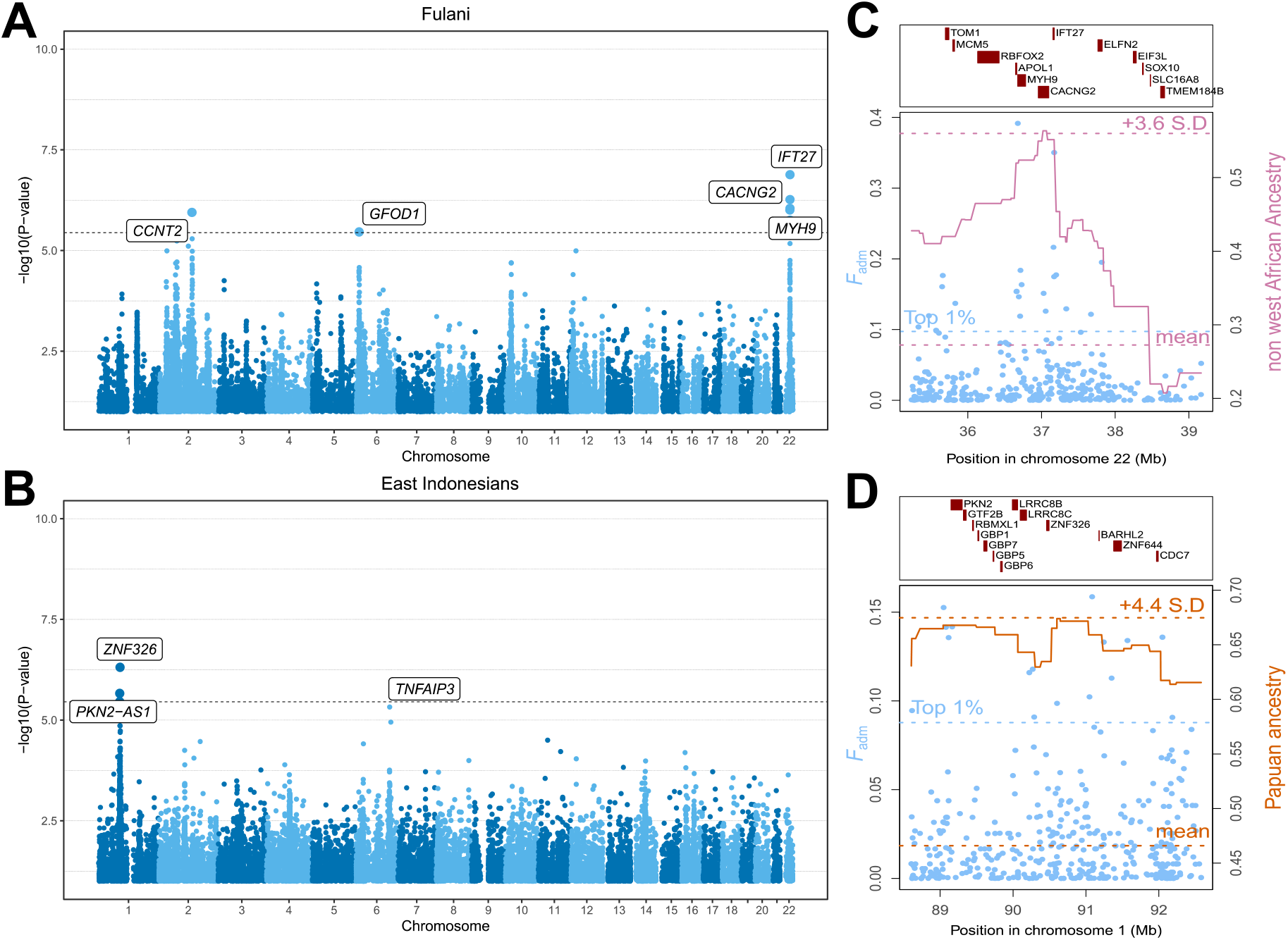
Newly discovered genomic signals of adaptive admixture. Genome-wide signals of adaptive admixture in (A) the Fulani nomads of West Africa and (B) East Indonesians. Highlighted blue points indicate variants that pass the Bonferroni significance threshold (shown by a horizontal dotted line). Gene labels were attributed based on the gene with the highest V2G score within 250-kb of the candidate variant. (C-D) Local adaptive admixture signatures for (C) the *IFT27/MYH9/APOL1* region in the Fulani nomads and (D) the *PKN2/LRR8CB* region in East Indonesians. Light blue points indicate *F*_adm_ values for individual variants. The pink and orange solid lines indicate the local ancestry from Europeans and North Africans, and Papuans, respectively.

### New candidate genes for adaptive admixture

We found several novel candidate loci for adaptive admixture (Fig 6 and S13 Fig), among which the *MYH9/APOL1* locus in the Fulani (Fig 6A and 6C; *P =* 1.3×10^−7^; top ranked SNP in *IFT27*; expected frequency of 0.15 *vs.* observed frequency of 0.45). Common *APOL1* variants confer both protection against human African trypanosomiasis (HAT, or sleep sickness) and susceptibility to common kidney diseases in African-descent individuals [60]. Another candidate is the *ZNF326*/*PKN2* locus in East Indonesians (*P =* 1.1×10^−6^; top ranked SNP in *ZNF326*; expected frequency of 0.27 *vs.* observed frequency of 0.46), which shows a large excess of Papuan ancestry (Fig 6B and 6D). *PKN2* plays a role in cellular signal transduction responses and has been reported as involved in the regulation of glucose metabolism in skeletal muscle [57]. A nearby locus, *LRRC8B*, has been reported as a candidate for positive selection in Solomon Islanders [41], although it did not show signals for adaptive admixture in this population. Nevertheless, a unique, strong signal was detected at the *ARRDC4*/*IGF1R* locus in Solomon islanders (*P =* 7.4×10^−9^; top ranked SNP close to *ARRDC4*; expected frequency of 0.91 *vs.* observed frequency of 0.42), where an excess of East Asian-related ancestry was observed (S13B and S13F Fig). This locus was previously identified as a candidate for positive selection in Near and western Remote Oceanians [41]. ARRDC4 is an arrestin that plays important roles in glucose metabolism and immune response to enterovirus infection [58], whereas IGF1R, the receptor for the insulin-like growth factor, is a key determinant of body size and growth [59,60]. A last example is *CXCL13* in the Nama pastoralists from South Africa (S13A and S13E Fig; *P =* 2.3×10^−6^; top ranked SNP identified in *CXCL13*; expected frequency of 0.49 *vs.* observed frequency of 0.20). The *CNOT6L/CXCL13* locus has previously been reported as suggestively associated tuberculosis (TB) risk in South African populations with San ancestry [61]. However, we found that the top-ranking variants show outlier extended haplotype homozygosity in the Ju|’hoansi San, used as source population (iHS = −3.12), while European ancestry is in excess at the locus in the Nama, suggesting a spurious signal due to positive selection in the proxy source population (S4 Fig).

We also detected suggestive signals of adaptive admixture at genes shown to be strong candidates for positive selection, including the *MCM6*/*LCT* locus in the Bantu-speaking Bakiga of Uganda (S14 Fig; *P =* 4.3×10^−6^; top ranked SNP in *CCNT2*; expected frequency of 0.15 *vs.* observed frequency of 0.31) and *TNFAIP3* in East Indonesians, who show an excess of Papuan-related ancestry at the locus (Fig 6A; *P =* 5.0×10^−6^; top ranked SNP in *TNFAIP3*; expected frequency of 0.27 *vs.* observed frequency of 0.43). The *TNFAIP3* locus has not only been reported as under positive selection in Papuan populations [41], but also as adaptively introgressed from Denisovans [41,62–64]. TNFAIP3 plays an important role in human immune tolerance to pathogen infections [65]. Collectively, these results indicate that adaptive admixture has occurred in various human admixed populations, and highlight metabolism and microbial infections as important drivers of recent genetic adaptation.

## Discussion

In this study, we evaluated the power of several neutrality statistics to detect loci under positive selection in admixed populations and used these statistics to explore cases of adaptive admixture in the genomes of 15 worldwide human populations. *F*_adm_ and LAD, or closely related statistics based on the difference between the observed and expected allele frequencies and admixture proportions, have been used in several empirical studies but their power has not been thoroughly evaluated. Here, we showed that these statistics are powerful to detect adaptive admixture and have no power to detect residual signals of positive selection in the source populations. Thus, *F*_adm_ and LAD are suited to scan genomes of admixed populations to search for loci under positive selection since admixture, particularly when selection is strong (i.e., *s* ≥ 0.05), admixture is relatively old (i.e., *T*_adm_ > 2,000 years) and the admixture proportion is moderate-to-low (i.e., *α <* 0.6). Notably, we found that power is marginally affected when admixture has been recurrent, a feature that is convenient given the difficulty to distinguish single-pulse, double-pulse or more complex admixture models from the genetic data [8,41–46]. Furthermore, we observed that *F*_adm_ is more powerful than LAD when selection occurs in the admixed population only, whereas LAD is more powerful than *F*_adm_ when source sample sizes are low (i.e., *n* = 20) and when the true and proxy source populations are distantly related (i.e., *F*_ST_ ≥ 0.02). The latter result is consistent with the known robustness of LAI to cases when the populations used as reference sources are poor proxies of the true source populations [39]. Nonetheless, caution must be taken when handling population proxies, as selection occurring in the proxy population only can produce genomic signals, for both LAD and *F*_adm_, that might be misinterpreted as adaptive admixture [21,22,41]. We suggest that performing selection scans on the proxy source populations can help distinguish false from true adaptive admixture signals. Finally, we estimated that combining *F*_adm_ and LAD statistics into a unified statistic, based on the Fisher’s method, confers higher power than when using them individually.

When applying this method on the empirical data, we identified several previously reported candidate variants for adaptive admixture. These include the *ACKR1* Duffy-null allele detected in admixed populations from Madagascar [20], the Sahel [25] and Pakistan [19], the lactase persistence −13910 C>T *LCT* allele in the Fulani from West Africa [56], and *HLA* alleles in Bantu-speaking population from western central Africa [21]. These candidate loci were initially detected based on LAD only, or in combination with classic neutrality statistics. However, the detection of natural selection with the LAD statistic has previously been questioned, because deviations in local ancestry can be explained as artefacts of long-range linkage disequilibrium (LD), which was not properly modelled by the first-generation LAI methods [66]. Our analyses reveal that these candidate genomic regions not only show outlier LAD values, but also outlier *F*_adm_ values. Because *F*_adm_ only depends on allele frequencies at the SNP of interest, our results support the view that the observed signals of adaptive admixture are genuine and unlikely to be explained by incorrectly modelled LD.

Our results also highlight novel signals of adaptive admixture, such as the *APOL1*/*MYH9* locus in the Fulani nomads of West Africa. Interestingly, an *APOL1* haplotype of non-African origin, named G3, was shown to be under positive selection in the Fulani of Cameroon [67], in line with the excess of non-African ancestry that we detected at the locus in the Fulani from Burkina Faso. Nevertheless, the physiological effect of the G3 variants is still debated: experimental work suggests that the G3 haplotype has no lytic activity against *Trypanosoma* parasites and is not associated with increased susceptibility to common kidney diseases in African Americans [68]. Alternatively, the significant excess of non-African ancestry observed at the locus may be due to strong negative selection against HAT-resistance *APOL1* alleles (i.e., G1 and G2 haplotypes), in regions where the incidence of sleeping sickness is low, such as Burkina Faso [69]. As they do not confer a selective advantage in *Trypanosoma brucei*-free regions, the G1 and G2 haplotypes only strongly increase the risk for chronic kidney diseases [70] and thus become disadvantageous. Further epidemiological and experimental work will be needed to confirm this hypothesis.

Several of the putatively selected alleles detected here have previously been shown to be under strong positive selection in humans, such as *ACRK1* [71–73], *LCT* [74,75] or *HLA* [76]. This is in accordance with our simulation study, which shows that, over the explored time period of five millennia, the strength of selection for the beneficial allele must be sufficiently high to leave detectable signatures in the genomes of admixed individuals. In addition to their confirmatory nature, these results improve our understanding of the selective advantage conferred by the detected beneficial alleles. Because *F*_adm_ and LAD detect natural selection since admixture only, selection studies in recently admixed populations represent a valuable tool to detect recent ongoing selection. Furthermore, admixed and source populations have often lived in different environments, so evolutionary studies of adaptive admixture can help refine correlations between signatures of natural selection and environmental pressures. An illustrative example is the Duffy-null *FY*B*^*ES*^ allele, which is fixed or nearly fixed in most sub-Saharan African populations [73]. It has long been proposed that natural selection has favoured this allele because it protects against malaria due to *Plasmodium vivax* [77]. Indeed, cellular experiments have shown that the parasite depends on the Duffy protein for erythrocytic infection [31,32]. However, recent studies have casted doubt on this result, because *P. vivax* has been detected in *FY*B*^*ES*^ homozygous carriers [78,79], suggesting that parasite invasion is possible when its human receptor ACKR1 is absent. We and others have found signatures of adaptive admixture for the *FY*B*^*ES*^ allele in African-descent admixed populations from Madagascar [12,20], Capo Verde [23], the Sahel [23], and Pakistan [19], but not in North Americans or South Africans [21,22]. Evidence of ongoing positive selection for Duffy negativity is thus confined to regions where the current incidence of *P. vivax* malaria is estimated to be high [80]. These findings thus support the view that resistance to *vivax* malaria is the main evolutionary force driving the frequency of the Duffy-null allele in humans.

Overall, our study reports evidence that recent admixture has facilitated human genetic adaptation to varying environmental conditions. It has been proposed that gene flow can promote rapid evolution when the demographic structure of a species is unstable [81]. Our findings support this view, as *Homo sapiens* is a structured species that has settled a large variety of ecotypes and has undergone large-scale, massive dispersals followed by extensive gene flow [7]. We thus anticipate that more cases of adaptive admixture in humans will soon be uncovered, thanks to methodological and technological advances. Importantly, given the highly conservative nature of our approach, it is very likely that we do not recover variants that have probably been weakly to mildly selected since admixture, such as *TNFAIP3* in Indonesian populations of Papuan-related ancestry [41,62–64] or the *MCM6*/*LCT* locus in the Bantu-speaking Bakiga from Uganda [21]. The development of new powerful neutrality statistics, such as the integrated decay in ancestry tracts iDAT [23], combined with model-based probabilistic frameworks [82], is a promising path to improve the power to detect adaptive admixture and better account for the demography of admixed populations. Furthermore, many human traits are known to be highly polygenic, suggesting that polygenic adaptation is a key driver of phenotypic evolution [83]. Thus, new methods are also required to detect polygenic selection since admixture [84]. Finally, genomic studies of adaptive admixture are expected to be more powerful when admixture is ancient, but statistical tests for admixture in modern genomes have low power when admixture time is older than 5,000 years [8]. Ancient genomics studies offer a great opportunity to circumvent this limitation, by revealing how human populations interacted in the past and how beneficial alleles have spread in time and space [33,85]..

## Methods

### General simulation settings

All the simulations were computed with the SLiM 3.2 engine [86] under the Wright-Fisher model. Each simulation consisted of a 2-Mb long locus characterized by varying recombination and mutation rates. For each simulation, we sampled the physical coordinates of a random 2 Mb genomic window in the human genome, excluding telomeric and centromeric regions, and assigned recombination rates based on the 1000 Genomes phase 3 genetic map [87] and mutation rates based on Francioli et al. mutation map [88]. We additionally incorporated background selection by simulating exon-like genetic elements positioned according to the position of exons in the sampled 2-Mb genomic window. Each simulated exon was made of positions under negative selection or under neutrality, mimicking non-synonymous and synonymous positions, respectively. Selection coefficients of negatively selected mutations were sampled from the European-based gamma distribution described in Boyko et al. 2008 [89]. Deleterious mutations were set to occur three times more frequently than neutral mutations, to account for codon degeneracy. Because we used computationally intensive forward-in-time simulations, we rescaled effective population sizes and times according to *N*/*λ* and *t*/*λ*, with *λ* = 10, and used rescaled mutation, recombination and selection parameters, *λμ*, *λr* and *λs* [86].

### Admixture with selection models

In the three simulated scenarios of admixture with selection, the following parameters were given fixed values: effective population sizes (source and admixed populations) *N*_e_ = 10,000; divergence time between source populations *T*_div_ = 2,000 generations; admixture proportion from the minor source *α =* 0.35; admixture proportion from the major source 1 – *α =* 0.65; and time of the single pulse admixture event *T*_adm_ = 70 generations.

For scenarios 1 and 2, the selected mutation was set to appear 350 generations ago in the source population that contributes the least to the admixed population and is transmitted to the admixed population with either the same selection coefficient (scenario 1) or a selection coefficient set to 0 (scenario 2). For scenario 3, we adapted a combination of recipes 9.6.2 and 14.7 from the SLiM manual [90] introducing a set of “ancestry marker” neutral mutations in the source population that contributes the least to the admixed population, and randomly choosing one of them to become beneficial by setting its selection coefficient to *s >* 0 in the admixed population only. We used 5 different values for the selection coefficient of the beneficial mutation ranging from *s* = 0.01 to *s* = 0.05. We computed 500 simulations for each admixture-with-selection scenario, as well as 500 simulations for the null scenario (no positive selection).

### Power of explored neutrality statistics

Neutrality statistics were computed for all polymorphisms within the 2-Mb simulated loci under no positive selection (H0), and only for the selected mutation for simulated 2-Mb loci under positive selection (H1). We estimated statistical power (i.e., the TPR) for each statistic as the proportion of values under H1 that are above a given H0 threshold (i.e., the FPR). We computed *F*_ST_, ΔDAF and iHS using selink [41]. We computed *F*_ST_ and ΔDAF between the admixed population and the source population that does not experience positive selection (in the case of scenario 3, where there is no selection in either source population, we used the source population with the major contribution). For iHS, we used a window of 200kb and normalized the values by bins of similar derived allele frequency (DAF).

For the admixture-specific statistics, we introduced an allele frequency-based statistic, *F*_adm_, that measures the difference between the observed and expected allele frequencies under admixture. *F*_adm_ is defined as follows:

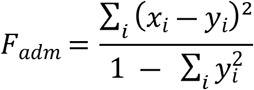

where *x_i_* and *y_i_* are the observed and expected frequencies of allele *i* in the admixed population, respectively, with *y_i_* = Σ_*p*_ *α_p_ x_p_* [37], i.e., the average of allele frequencies *x_p_* observed in the source populations *p* weighted by estimated admixture proportions *α_p_*. When calculating *F*_adm_ in simulated data, we used the simulated admixture proportions. Additionally, in both simulated and observed data, we excluded sites where the observed allele frequency in the admixed population *x_i_* is higher (or lower) than the maximum (or minimum) of the frequencies *x_p_* in the source populations. Although this can reduce the detection power in scenario 3, this filter increases power for adaptive admixture scenario (S2 Fig), which is the focus of this study.

We also computed a LAI based neutrality statistic, LAD, which measures the local ancestry deviation from the average genome-wide ancestry, defined as follows:

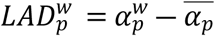

where, for a given window *w*, *α_p_* is the average local ancestry across admixed individuals and 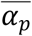 is the genome-wide admixture proportion. When calculating LAD with simulated data, we used the estimated average ancestry across all simulations [10,11,27]. We used RFMix v1.5.4 [39] to estimate local ancestry, with default parameter values (except for –G, which was replaced with the simulated *T*_adm_ value) and using the forward-backward option with 3 expectation maximization steps. Because LAD is sensitive to phasing errors [39], we incorporated potential phasing errors in our simulations by phasing with SHAPEIT v4.2.1 [91] unphased, simulated diploid individuals obtained from the combination of two haploid individuals.

### Sample size and source population choice scenarios

We explored 5 different values of sample sizes for each of the source populations and the admixed population: *n* = 20, 50, 100, 200 and 500 individuals. When exploring the values for a given population, sample sizes for the other two were fixed to *n* = 50 individuals. For the source population choice, we explored 4 different genetic distance values (in a *F*_ST_ scale) between the proxy population and the true source population. The explored *F*_ST_ values were the following: 0.005, 0.01, 0.02 and 0.03. These values were obtained by fixing the divergence time between the true source population and the proxy to 400 generations and setting the effective population size of the proxy to 10,000, 4000, 1000 and 500 respectively. For the scenario of selection on the parental proxy only, we randomly selected a mutation present in both proxy and true source population 1 and assigned it a selection coefficient of *s* = 0.02. To compare the excess of source population 2 ancestry generated by this scenario, we additionally simulated an adaptive admixture scenario where the beneficial mutation was transmitted to the admixed population from source population 2. In all these scenarios, the following parameters were given a fixed value: *N*_e_ = 10,000; *T*_div_ = 2,000 generations; *α =* 35%; 1 – *α =* 65%; *T*_adm_ = 70 generations and *s* = 0.02.

### Non-stationary demography

We simulated four alternative demographic scenarios, each with 500 simulations under adaptive admixture and 500 simulations with no positive selection, to estimate detection power. In all six scenarios the following parameters were given a fixed value: *T*_div_ = 2,000 generations; *α =* 35%; 1 – *α =* 65%; *T*_adm_ = 70 generations and *s* = 0.02. Demographic scenarios include (i) a recent expansion of the source population, where the source population (*N*_e_ = 10,000) undergoes an expansion with a 5% growth rate since *T*_adm_; (ii) a recent expansion of the admixed population, where the admixed population (*N*_e_ = 10,000) undergoes an expansion with a 5% growth rate since *T*_adm_; an old expansion of the source population, where the source population (*N*_e_ = 10,000) undergoes an expansion with a 5% growth rate since *T*_adm_ + 500 generations; (iii) an old bottleneck in the source population, where the source population (*N*_e_ = 10,000) undergoes a 10-fold *N*_e_ reduction from *T*_div_ to *T*_div_ – 50; and (iv) a recent bottleneck in the admixed population, where the admixed population (*N*_e_ = 10,000) undergoes a 10-fold *N*_e_ reduction from *T*_adm_ to *T*_adm_ – 50. We compared these scenarios to a constant population size scenario, with the same general parameters and the *N*_e_ of all populations fixed to 10,000.

### Complex admixture scenarios

We estimated detection power under 2 additional admixture scenarios. Under the double pulse model, the admixed population diverges from one of the source populations and receives two separate admixture pulses from the other source population, of the same intensity. Under the constant continuous admixture, the admixed population diverges from one of the source populations and receives admixture pulses at each generation from the source population, of constant intensity. For these scenarios to be comparable to the single pulse admixture scenario, we set the sum of the admixture proportions contributed by each pulse to be equal to *α =* 35%, and the average of the admixture dates to be equal to 70 generations. Namely, for the double pulse scenario, each pulse contributed *α =* 17.5% and took place 130 and 10 generations in the past, and for the constant continuous scenario, the first instance of admixture took place 130 generations in the past, and each instance contributed approximately *α =* 35% / 130 = 0.27%, when *T*_adm_ is not rescaled, and *α =* 2.7%, when *T*_adm_ is rescaled.

### Admixture parameters

Under the single pulse admixture model, we explored detection power as a function of different model parameters, which are shown in S1 Table. In total, over 20,000 unique parameter combinations were explored, reason by which the number of simulations was reduced from 500 to 100, to limit computational burden. For the frequency of the beneficial mutation on the source population at the time of admixture, instead of conditioning on the frequency within simulations (which would have drastically increased the computational intensity), we introduced the beneficial mutation *T*_mut_ generations ago, in the source population, based on previous results [92]. For each statistic and each combination, we calculated the proportion of simulated sites under selection that were recovered using a threshold of FPR = 5%. We then averaged the power across demographic parameter values to obtain a single value for each combination of *T*_adm_, *α* and *s*. We performed a similar procedure to obtain a single value for each combination of *T*_adm_, *α*, and one of the other parameters (S6-10 Figs).

### Empirical detection of adaptive admixture

We analysed the genomes of 15 admixed populations to find signals of adaptive admixture. The datasets and references for all admixed and source populations can be found in S2 Table, as well as the final SNP count after merging admixed and source population datasets. For each merged dataset, we: (i) excluded sites with a proportion of missing genotypes > 5%, using plink 2.0 [93]; (ii) excluded A/T and C/G variant sites; (iii) excluded first and second degree-related individuals (kinship coefficient > 0.08 computed with KING v2.2.2 [94]) and (iv) performed phasing using SHAPEIT v4.2.1, using default parameter values. Additionally, we verified the validity of an admixture model for each set of source/admixed populations by computing admixture *f3* statistics with admixr package version 0.7.1[95] (S2 Table). Admixture proportions were obtained by running ADMIXTURE v1.23 [96], considering the *K* value producing the lowest cross-validation error and a set of “independent” SNPs obtained by running the ‘--indep-pairwise’ command with plink 2.0, with the following parameters: 50-SNP window, 5-SNP step, and *r*^2^ threshold of 0.5. We also verified that the chosen value for *K* matched the number of source populations for the studied admixed population. Local ancestry was obtained with RFMix v.1.5.4 [39], after excluding 2 Mb at telomeres and centromeres of each chromosome, as well as monomorphic sites and singletons, and using default parameter values except for the generation time ‘-G’, which was given an estimated value for each population based on literature (S2 Table).

In addition of *F*_adm_ and LAD, we combined the SNP ranks of these two statistics using Fisher’s method, defined as follows:

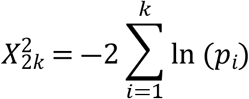

where *p_i_* is defined as the rank of a given SNP for the statistic *i*, divided by the total number of analysed SNPs (i.e., the empirical *P*-value), and *k* = 2 is the number of statistics. Using simulations, we verified that this statistic followed a chi-squared distribution with 2*k* = 4 degrees of freedom under no positive selection, even when the admixed population experienced a 10-fold bottleneck (Fig 4A). In these simulations, we used the same parameter values as those in Fig 2C for the “constant size” and “bottleneck in the admixed population” scenarios. Statistical significance was defined based on Bonferroni correction: we considered a *P*-value threshold of 0.05 divided by the number of effective 0.2-cM RFMix windows analysed (all SNPs within the same window had the same local ancestry value), which yielded on average, a *P*-value of 3.5×10^−6^ threshold (S2 Table). To annotate the different signals that passed this threshold, we chose the protein coding gene within 250-kb of the variant with the highest V2G score [97].

## Data Availability Statement

Accession numbers for the SNP array data used in this study are listed in S2 Table. All SLiM parameter files can be found here: https://github.com/h-e-g/ADAD.

## Acknowledgements

We thank all volunteers participating in this research; Sophie Créno and the HPC Core Facility of Institut Pasteur (Paris) for the management of computational resources; Omar Alva Sanchez, Denis Pierron, Thierry Letellier, Mario Vicente, Carina Schlebusch, Andres Moreno-Estrada, Andres Ruiz-Linares and the Health Aging and Body Composition (Health ABC) Study for kindly providing access to their data. We also thank Javier Bougeard, Lara Rubio Arauna, Jérémy Choin, Maxime Rotival, Paul Verdu and Olivier Tenaillon for helpful discussions. S.C.-E. is supported by Sorbonne Université Doctoral College, the Inception program (Investissement d’Avenir grant ANR-16-CONV-0005) and the Institut Pasteur. The laboratory of Human Evolutionary Genetics is supported by the Institut Pasteur, the Collège de France, the CNRS, the Fondation Allianz-Institut de France, the French Government’s Investissement d’Avenir programme, Laboratoires d’Excellence ‘Integrative Biology of Emerging Infectious Diseases’ (ANR-10-LABX-62-IBEID) and ‘Milieu Intérieur’ (ANR-10-LABX-69-01), the Fondation de France (n°00106080), and the Fondation pour la Recherche Médicale (Equipe FRM DEQ20180339214) and the French National Research Agency (ANR-19-CE35-0005).

## Author contributions

E.P. conceived and supervised the project. S.C.-E. designed and performed all the analyses, with critical input from G.L. G.L. provided theoretical and methodological context. S.C.-E. and E.P. wrote the manuscript, with critical input from G.L. and L.Q.-M.

## Supporting Information

**S1 Fig. Performance of neutrality statistics under different scenarios of admixture with selection and selection coefficients.** Receiver operating characteristic (ROC) curves comparing classic neutrality statistics *F_ST_*, ΔDAF and iHS and admixture specific statistics *F*_adm_ and LAD, across the 3 explored admixture with selection scenarios, with varying selection coefficients.

**S2 Fig. Performance of *F*_adm_ when applying or not an allele frequency filter, under different selection with admixture scenarios.** Receiver operating characteristic (ROC) curves comparing *F*_adm_, with and without applying an allele frequency filter based on the source populations (see Methods), under the 3 explored admixture with selection scenarios.

**S3 Fig. Effects of sample size on *F*_adm_ and LAD.** (A) Distributions under the null hypothesis (no positive selection) of *F*_adm_ and LAD, with varying sample sizes for the admixed population. (B) Effect of the sample size of the source populations on the detection power of *F*_adm_ and LAD.

**S4 Fig. False positive signals due to selection in the proxy source population.** (A) Observed *vs.* expected allele frequencies in the admixed population, when there is or not positive selection in the proxy source population. (B) Distributions of local ancestry in the admixed population from the unselected source population, when there is or not positive selection in the proxy source population. (C) ROC curves for *F*_adm_ and LAD comparing the scenario where there is positive selection in the proxy source population (in teal; proxy of the source population 1) and the scenario where there is a true, adaptive admixture event (in salmon). The beneficial mutation is selected in the source population that has no proxy (population 2). (D–E) Absolute iHS values in the proxy source population *vs.* (D) *F*_adm_ and LAD values for the selected mutation in the admixed population, when there is selection in this proxy, or where there is adaptive admixture coming from the source population with no proxy (population 2). Colour codes are the same as for panel C. Dashed green lines represent the 99^th^ percentiles (based on the null model simulations) for absolute iHS (vertical) and *F*_adm_ or LAD (horizontal). Excluding values that are above the absolute iHS 99^th^ percentile excludes approximately 90% of the extreme *F*_adm_ and LAD values generated by the selection on proxy scenario but, importantly, does not exclude any extreme value generated by the true adaptive admixture scenario.

**S5 Fig. Effects of complex admixture and non-stationary demography on the power to detect adaptive admixture.** (A) *F*_adm_ and LAD detection power for a FPR = 5% in different admixture scenarios: a single pulse scenario, a double pulse scenario and a constant continuous admixture scenario (Methods). (B) Distributions of *F*_adm_ and LAD under the null hypothesis (no positive selection), with or without a 10-fold bottleneck in the admixed population.

**S6 Fig. Effects of the divergence time between source populations on the power to detect adaptive admixture.** Effects on the detection power of *F*_adm_ and LAD of admixture time *T*_adm_, admixture rate *α* and *T*_div_. Colour indicates average detection power for a FPR = 5% threshold, across combinations of the remaining parameters.

**S7 Fig. Effects of effective population sizes on the power to detect adaptive admixture.** Effects on the detection power of *F*_adm_ and LAD of admixture time *T*_adm_, admixture rate *α* and *N*_1_, *N*_2_ and *N*_adm_, the effective population sizes of source population 1, source population 2 and the admixed population, respectively. Colour indicates average detection power for a FPR = 5% threshold, across combinations of the remaining parameters.

**S8 Fig. Effects of the frequency of the beneficial mutation (*s =* 0.01) on the power to detect adaptive admixture.** Effects on the detection power of *F*_adm_ and LAD of admixture time *T*_adm_, admixture rate *α* and *F*_onset_, the frequency of the beneficial mutation in the source population at the time of admixture *T*_adm_. Colour indicates average detection power for a FPR = 5% threshold, across combinations of the remaining parameters.

**S9 Fig. Effects of the frequency of the beneficial mutation (*s =* 0.05) on the power to detect adaptive admixture.** Effects on the detection power of *F*_adm_ and LAD of admixture time *T*_adm_, admixture rate *α* and *F*_onset_, the frequency of the beneficial mutation in the source population at the time of admixture *T*_adm_. Colour indicates average detection power for a FPR = 5% threshold, across combinations of the remaining parameters.

**S10 Fig. Effects of the frequency of the beneficial mutation (*s =* 0.10) on the power to detect adaptive admixture.** Effects on the detection power of *F*_adm_ and LAD of admixture time *T*_adm_, admixture rate *α* and *F*_onset_, the frequency of the beneficial mutation in the source population at the time of admixture *T*_adm_. Colour indicates average detection power for a FPR = 5% threshold, across combinations of the remaining parameters.

**S11 Fig. Distributions of Fisher’s combined *P*-values in the empirical data.** Histograms of combined *P*-values using Fisher’s method, for the 15 analysed admixed populations. The *P*-values are uniformly distributed, except for certain populations where there is an excess of small *P*-values, corresponding to the populations where signals for adaptive admixture were found.

**S12 Fig. Other previously reported genomic signals of adaptive admixture.** Genome-wide signals of adaptive admixture in (A) Malagasy populations from Madagascar and (B) African-descent Makranis and Makrani Baluch from Pakistan. Highlighted blue points indicate variants that passed the Bonferroni significance threshold (shown by a horizontal dotted line). Gene labels were attributed based on the gene with the highest V2G score within 250-kb of the candidate variant. (C-D) Local adaptive admixture signatures for the *ACKR1* region in (C) Malagasy from Madagascar and (D) Makranis and Makrani Baluch from Pakistan. Light blue points indicate *F*_adm_ values for individual variants. The gold solid line indicates the average African local ancestry.

**S13 Fig. Other novel genomic signals of adaptive admixture.** Genome-wide signals of adaptive admixture in (A) the Nama from South Africa, (B) Solomon Islanders, (C) Vanuatu Islanders and (D) admixed Peruvians. Highlighted blue points indicate variants that passed the Bonferroni significance threshold (shown by a horizontal dotted line). Gene labels were attributed based on the gene with the highest V2G score within 250-kb of the candidate variant. (E-H) Local adaptive admixture signatures for (E) the *CNOT6L/CXCL13* region in the Nama from South Africa, (F) the *ARRDC4* region in Solomon Islanders, (G) the *IGKV1-17* region in Vanuatu Islanders and (H) the *ITPR2* region in admixed Peruvians. Light blue points indicate *F*_adm_ values for individual variants. The yellow, gold and pink solid lines indicate average local ancestry from East Africans, Austronesians and Europeans respectively.

**S14 Fig. Genome scans for populations where there is no evidence for adaptive admixture.** Manhattan plots of –log_10_(*P*-values) for the combined Fisher’s method, in the remaining 6 admixed populations where no variant passes the Bonferroni significance threshold (shown by a horizontal dotted line).

**S1 Table. Simulated parameter values for adaptive admixture models.**

**S2 Table. Studied populations.**

**S3 Table. Significance thresholds for genome scans of adaptive admixture.**

